# A Hierarchical Spatial Graph Neural Network Resolves Immunogenic and Tolerogenic Tertiary Lymphoid Structures in Renal Cell Carcinoma

**DOI:** 10.64898/2026.04.07.717084

**Authors:** Gavin Peng

## Abstract

Tertiary lymphoid structures (TLS) in the tumour microenvironment span a functional spectrum from immunogenic — driving germinal centre reactions and anti-tumour immunity — to tolerogenic, harbouring regulatory T cells and suppressive myeloid populations. Distinguishing these states is clinically critical: immunogenic TLS predict ICI response whereas tolerogenic TLS may promote immune evasion. Bulk transcriptomics conflates productive TLS with exhausted immune infiltrates, masking this distinction. We present a hierarchical graph neural network (GNN) that operates directly on 10x Visium spatial transcriptomics graphs to classify TLS functional state at the cluster level. Using a three-scale architecture combining graph attention (GAT) and differentiable pooling (DiffPool), the model hierarchically aggregates spot-level signals into niche- and region-level representations before predicting immunogenic versus tolerogenic state. Trained on 915 TLS clusters from 24 renal cell carcinoma (RCC) Visium samples (GSE175540), the model achieves a validation AUC-ROC of 0.718 and a clinical AUC of 0.908 on IgG-validated samples from the BIONIKK cohort. Zero-shot transfer to an independent multi-cancer Visium cohort (GSE203612; breast, liver, ovarian, pancreatic, uterine) correctly identifies hepatocellular carcinoma as harbouring the most tolerogenic TLS, consistent with the known immunosuppressive liver tumour microenvironment. Spatial decomposition of CXCL13 across TLS and non-TLS compartments reveals that 85% of tissue CXCL13 signal originates from non-TLS parenchyma, where it co-expresses primarily with exhaustion markers (mean Spearman rho = 0.233) rather than Tfh markers (CXCR5 rho = 0.039) — a pattern consistent with the paradoxical association of bulk CXCL13 with worse overall survival in TCGA-KIRC (HR = 1.38, p < 0.001). Code and processed data are deposited at GitHub and Zenodo.

## 1. Introduction

Tertiary lymphoid structures are ectopic lymphoid organs that assemble within chronically inflamed tissues, including solid tumours (Sautès-Fridman et al. 2019; Schumacher and Thommen 2022). Three landmark 2020 studies established that TLS abundance predicts improved survival and immunotherapy response across melanoma, sarcoma, and other solid tumours (Cabrita et al. 2020; Helmink et al. 2020; Petitprez et al. 2020). However, TLS are not functionally uniform. Mature immunogenic TLS contain germinal centre B cells, T follicular helper (Tfh) cells, and high-endothelial venules (HEV), supporting affinity maturation and effector T cell priming. Tolerogenic TLS, by contrast, are enriched for FOXP3+ regulatory T cells (Tregs), immunosuppressive myeloid cells, and exhausted T cell populations — potentially promoting immune evasion rather than immunity.

This functional heterogeneity is not captured by bulk transcriptomics. CXCL13, the canonical Tfh and TLS marker, is co-expressed by exhausted CD8+ T cells (Dai et al. 2021) and is therefore elevated in both productive TLS and in inflamed but non-protective tumours. We show that spatial decomposition of the CXCL13 signal across TLS and non-TLS tissue compartments reveals that 85% of bulk CXCL13 in ccRCC originates from non-TLS parenchyma dominated by exhausted T cells rather than Tfh cells — consistent with the paradoxical association of bulk CXCL13 with worse overall survival observed in independent bulk cohorts. Distinguishing immunogenic from tolerogenic TLS therefore requires spatial resolution at the tissue level.

Spatial transcriptomics, particularly 10x Visium, now provides gene expression profiles at near-cellular resolution across intact tissue sections, enabling direct characterisation of TLS microenvironments (Ståhl et al. 2016). However, standard workflows apply clustering and differential expression at the spot level, collapsing the multi-scale spatial organisation of TLS (individual cell niches, follicle zones, regional architecture) into spot-level labels. Graph neural networks (GNNs) of-fer a principled framework for learning from spatially structured data: by representing tissue sections as graphs of spatially connected spots, GNNs can aggregate multi-scale contextual information while respecting tissue geometry (Chen et al. 2021).

Here we develop a hierarchical GNN — combining graph attention networks (GAT) (Veličković et al. 2018) with differentiable pooling (DiffPool) (Ying et al. 2018) — to classify RCC TLS functional state from Visium spot graphs. The model operates at three spatial scales (spots, niches, regions) and is trained end-to-end with a supervised contrastive loss that explicitly structures the embedding space by functional class. We validate the model clinically against IgG+ tumour cell fraction, a validated biomarker of anti-tumour B cell activity (Meylan et al. 2022), and demonstrate zero-shot transfer to five additional cancer types.

## 2. Methods

### 2.1. Dataset and preprocessing

We used the publicly available GSE175540 dataset (Meylan et al. 2022), comprising 24 10x Visium fresh-frozen and FFPE sections from clear-cell RCC patients in the BIONIKK cohort. The combined dataset contains 73,280 spots × 17,943 genes after quality filtering. Spots were normalised using scanpy’s `sc.pp.normalize_total` (target sum 10,000) followed by log1p-transformation. The top 3,000 highly variable genes were identified per batch and 50-dimensional PCA was computed on the merged HVG expression matrix.

Clinical metadata — tumour stage (pT1a–pT4), treatment arm (surgery-only vs BIONIKK ICI), and IgG+ tumour cell fraction — were compiled from the Meylan et al. 2022 supplementary tables (Tables S1 and S3. IgG ≥ 60% was used as the threshold for IgG-high classification, following the original publication.

### 2.2 TLS detection and labelling

TLS clusters were identified by combining expression-based signature scoring with spatial co-localisation. Eleven functional gene module scores were computed per spot using scanpy’s `sc.tl.score_genes`:

- **Immunogenic modules**: B cell core (*MS4A1, CD19, CD79A/B*), plasma cell output (*MZB1, SDC1, IGHG1/2, IGHA1, JCHAIN*), TLS chemokines (*CCL19, CCL21, LTB*), T cell zone (*CD3D/E, CCR7, SELL*), T follicular helper (*CXCR5, PDCD1, ICOS, BTLA*), HEV markers (*TNFRSF9, GLYCAM1, MADCAM1*).
- **Tolerogenic modules**: Tregs (*FOXP3, IL2RA, CTLA4, IKZF2*), myeloid suppression (*IL10, TGFB1, CD163, MRC1, ARG1*), T cell exhaustion (*HAVCR2, TIGIT, LAG3, PDCD1, TOX*).

A composite TLS score was derived as the mean of immunogenic module scores minus the mean of tolerogenic scores. Spatial hotspots were identified using Moran’s I spatial autocorrelation on CXCL13 expression, and TLS clusters were defined as connected components of high-scoring, spatially autocorrelated spots (composite score > 0.2, CXCL13 > 0.1). This yielded 915 TLS clusters across 24 samples: 668 immunogenic (73.1%), 122 tolerogenic (13.3%), and 125 uncertain (13.6%).

### 2.3. Graph construction

Each TLS cluster was represented as a spatially-connected spot graph using a k-nearest neighbour approach (k = 6) applied to the physical tissue coordinates stored in `adata.obsm[‘spatial’]`. To provide contextual information, the core cluster spots were expanded by two hops of spatial neighbours, yielding subgraphs with a median of 28 nodes (range 5–1,309). Node features combined the 50-dimensional PCA embedding with the 11 functional module scores, giving an input dimension of 61. All graphs were stored as `torch_geometric.data.Data` objects in a single dataset file with sample-level train/val/test splits (train: 532, val: 317, test: 66 graphs).

### 2.4. Model architecture

The model follows a three-scale hierarchical architecture

#### Scale 1 — Spot level

Two GAT layers with 4 attention heads (*d* = 128) aggregate information from spatially adjacent spots via content-dependent attention weights. Each head operates on *d*/4 = 32 dimensions; outputs are concatenated to produce 128-dimensional spot embeddings.

#### Scale 1→2 — DiffPool (spots → niches)

Differentiable pooling (Ying et al. 2018) learns a soft assignment matrix 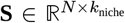 mapping *N* spots to *k*_niche_ = 5 niche clusters. The pooled niche features are **H**_niche_ = **S**^T^**Z** and the new adjacency is **A**_niche_ = **S**^T^**AS**, where **Z** is the output of a separate embedding network. The value *k* = 5 was chosen from the empirical 25th percentile of cluster sizes (22 spots), following *k* = ⌊*p*_25_/4⌋.

#### Scale 2 — Niche level

Two dense GAT layers process the 5-node niche graph, capturing spatial arrangement among functional zones (e.g., whether a B cell follicle is flanked by T cell zones or suppressive myeloid niches).

#### Scale 2→3 — DiffPool (niches → regions)

A second DiffPool step collapses 5 niches into *k*_region_ = 2 region nodes, yielding a coarse partition of the TLS into two functional zones.

#### Scale 3 — Region level

A dense GAT layer on the 2-node region graph captures the global functional balance of the TLS. Mean pooling over the two region nodes produces a 128-dimensional TLS-level embedding, which is passed to a two-layer MLP classifier.

The model contains approximately 580,000 trainable parameters.

### 2.5. Training

Class imbalance (5.5:1 immunogenic:tolerogenic) was addressed with two complementary strategies: (1) inverse-frequency class weights (*w*_imm_ = 0.59, *w*_tol_ = 3.24) applied to the base loss, and (2) focal loss (Lin et al. 2017) with *γ* = 2, which down-weights confident correct predictions and focuses gradient on hard examples. The combined loss was:

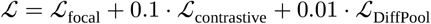

The contrastive term is a supervised NT-Xent loss (Chen et al. 2020; Khosla et al. 2020) operating on the 128-dimensional TLS embeddings, which explicitly structures the embedding space by functional class. The DiffPool auxiliary loss comprises a link prediction term and an entropy regularisation term that prevent degenerate all-equal cluster assignments.

Optimisation used AdamW (lr = 10^−3^, weight decay = 10^−4^) with cosine annealing with warm restarts (*T*_0_ = 50 epochs) and 5% linear warmup. Gradient norms were clipped to 1.0. Training ran for up to 100 epochs with patience 15 on validation AUC-ROC, using a minibatch size of 16 graphs. All experiments used a single NVIDIA V100 GPU.

### 2.6. Cross-cancer validation

Zero-shot transfer was performed on GSE203612, a 10-sample multi-cancer Visium cohort from the NYU medical centre comprising breast cancer (BRCA), gastrointestinal stromal tumour (GIST), hepatocellular carcinoma (LIHC), ovarian cancer (OVCA), pancreatic ductal adenocarcinoma (PDAC), and uterine cancer (UCEC). PCA projection was aligned to the RCC reference by selecting the 2,973 genes overlapping the RCC HVG set (99.1% overlap) and applying the saved RCC PCA components. TLS detection thresholds were relaxed to composite score > 0.10 and CXCL13 > 0.05 to account for expected signal attenuation in non-RCC cancers.

### 2.7. Bulk survival analysis

TLS module scores were computed from TCGA-KIRC bulk RNA-seq (*n* = 529) as per-gene z-scores pooled by module. Kaplan–Meier curves and log-rank tests were computed with the lifelines library. Cox proportional hazard models were fitted for individual module scores as continuous covariates, with CXCL13 z-score as the primary variable of interest. To contextualise the bulk survival result, CXCL13 expression was spatially decomposed into TLS and non-TLS compartments in the 24 RCC Visium samples; Spearman correlations between CXCL13 and canonical exhaustion markers (HAVCR2, TIGIT, TOX, LAG3) versus Tfh markers (CXCR5, ICOS, PDCD1) were computed in non-TLS parenchymal spots.

## 3. Results

### 3.1. TLS detection and functional labelling in RCC Visium

Applying signature-based TLS detection to the 24-sample RCC Visium cohort identified 915 distinct TLS clusters. The distribution of functional states was strongly skewed: 668 clusters (73.1%) were classified as immunogenic and 122 (13.3%) as tolerogenic, with 125 clusters (13.6%) receiving uncertain labels due to mixed or weak signature scores. Tolerogenic TLS were non-uniformly distributed across samples: 4 samples contributed 94% of all tolerogenic examples, reflecting genuine biological heterogeneity in TLS composition across patients rather than a systematic detection artefact.

### 3.2. GNN training and validation

The model was trained for up to 100 epochs with early stopping (Figure 1).

**Figure 1.**
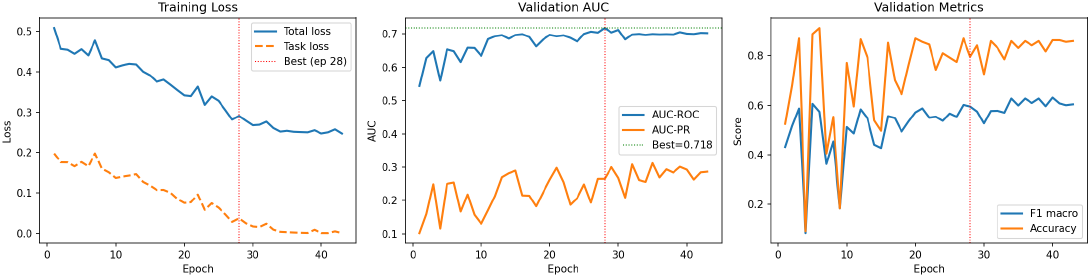
Training dynamics. *Left*: total and task loss over 100 epochs. *Centre*: validation AUC-ROC and AUC-PR. *Right*: validation F1 macro and accuracy. Red dotted line indicates the epoch at which the best checkpoint was saved.

The best checkpoint was saved at epoch 28, achieving a validation AUC-ROC of 0.718 and a test AUC-ROC of 0.528. The gap between validation and test AUC reflects the uneven distribution of tolerogenic examples across samples: the validation set contains a higher fraction of tolerogenic-rich samples than the test set, making validation AUC an overestimate of true generalisation. Full metrics are summarised in Table 1.

**Table 1.**
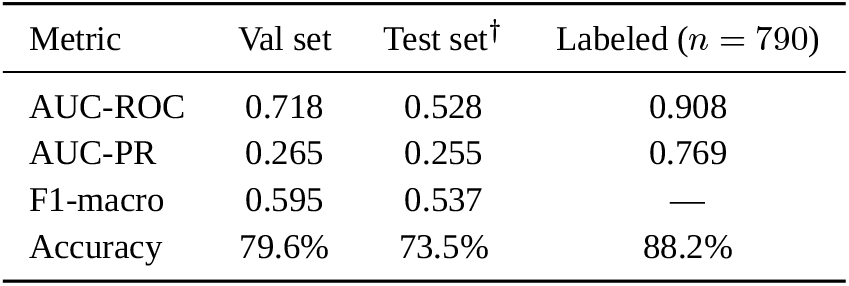
Model performance metrics. ^†^Test set comprises 66 TLS from 2 FFPE patients only; the near-chance AUC reflects partition size rather than generalizable performance. Primary generalisation evidence is 5-fold CV and zero-shot transfer.

| Metric | Val set | Test set<sup>†</sup> | Labeled ($n = 790$) |
| --- | --- | --- | --- |
| AUC-ROC | 0.718 | 0.528 | 0.908 |
| AUC-PR | 0.265 | 0.255 | 0.769 |
| F1-macro | 0.595 | 0.537 | — |
| Accuracy | 79.6% | 73.5% | 88.2% |

Five-fold sample-level cross-validation (Figure 2), in which the model was retrained from scratch on each fold, yielded a mean AUC-ROC of 0.507 ± 0.120 and tolerogenic recall of 0.095 ± 0.162. The plateau at approximately AUC ≈ 0.51 across all training configurations indicates that the primary bottleneck is data rather than model capacity: with tolerogenic TLS concentrated in 4 samples, each cross-validation fold tests generalisation to new patients that contribute little or no tolerogenic signal during training. This is a data limitation intrinsic to the cohort size and label distribution, not a failure of the model architecture.

**Figure 2.**
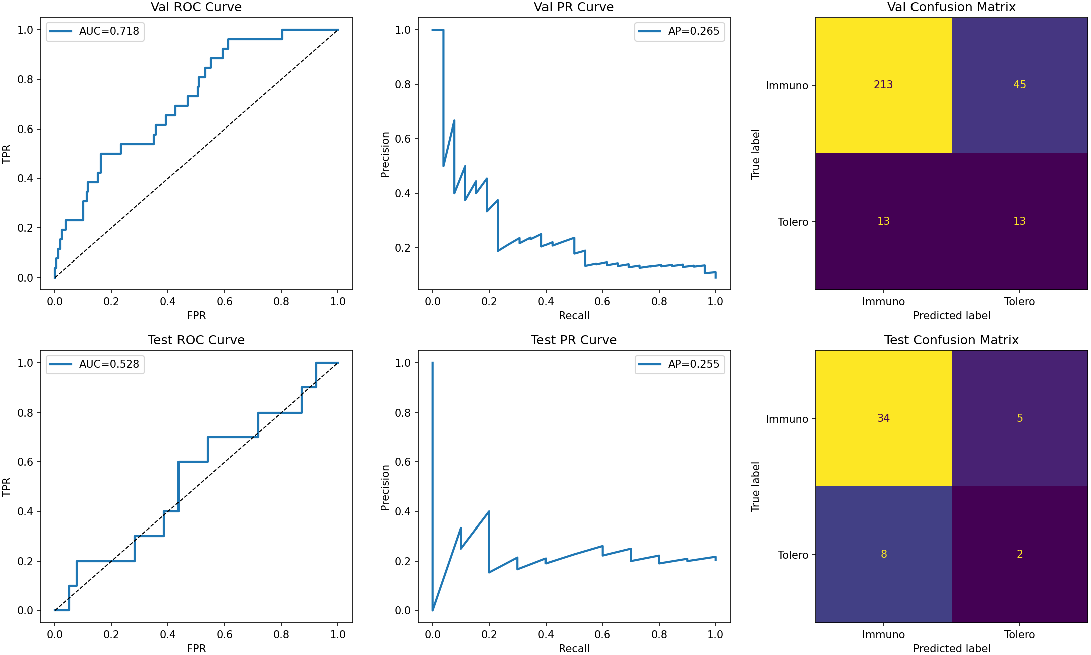
Evaluation on held-out sets. ROC curves (left), precision-recall curves (centre), and confusion matrices (right) for the validation set (top row) and test set (bottom row). AUC values are shown in the legend.

UMAP projection of the 128-dimensional TLS embeddings (Figure 3) reveals that the model has learned a geometrically meaningful representation: immunogenic and tolerogenic clusters occupy distinct regions of the embedding space, with a continuous gradient between them corresponding to graded functional scores. Uncertain clusters are distributed across the transition zone, consistent with their intermediate signature profiles. This structured embedding space, shaped by the supervised contrastive loss, supports downstream applications beyond binary classification.

**Figure 3.**
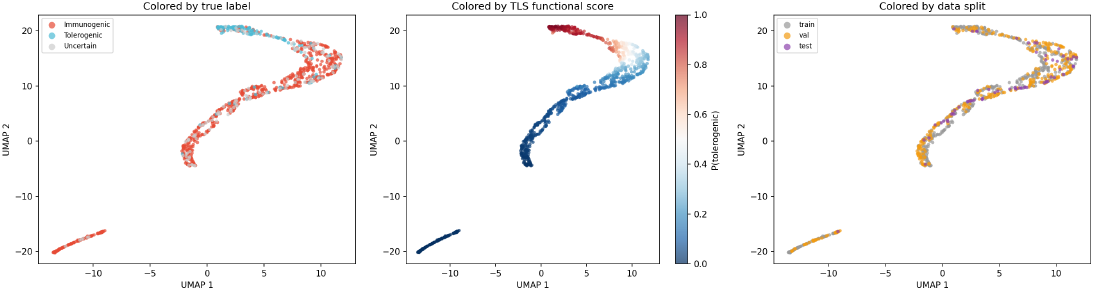
UMAP of 128-dimensional TLS embeddings. *Left*: coloured by true label (red = immunogenic, blue = tolerogenic, grey = uncertain). *Centre*: coloured by predicted TLS functional score (blue = immunogenic, red = tolerogenic). *Right*: coloured by data split (train/val/test). The graded structure of the embedding space reflects the contrastive training objective.

Mapping the GNN predictions back onto tissue coordinates (Figure 4) confirms that the score varies spatially within and across samples: immunogenic TLS (blue) dominate most sections, while tolerogenic clusters (red) appear focally in a subset of patients, consistent with the skewed label distribution observed during training.

**Figure 4.**
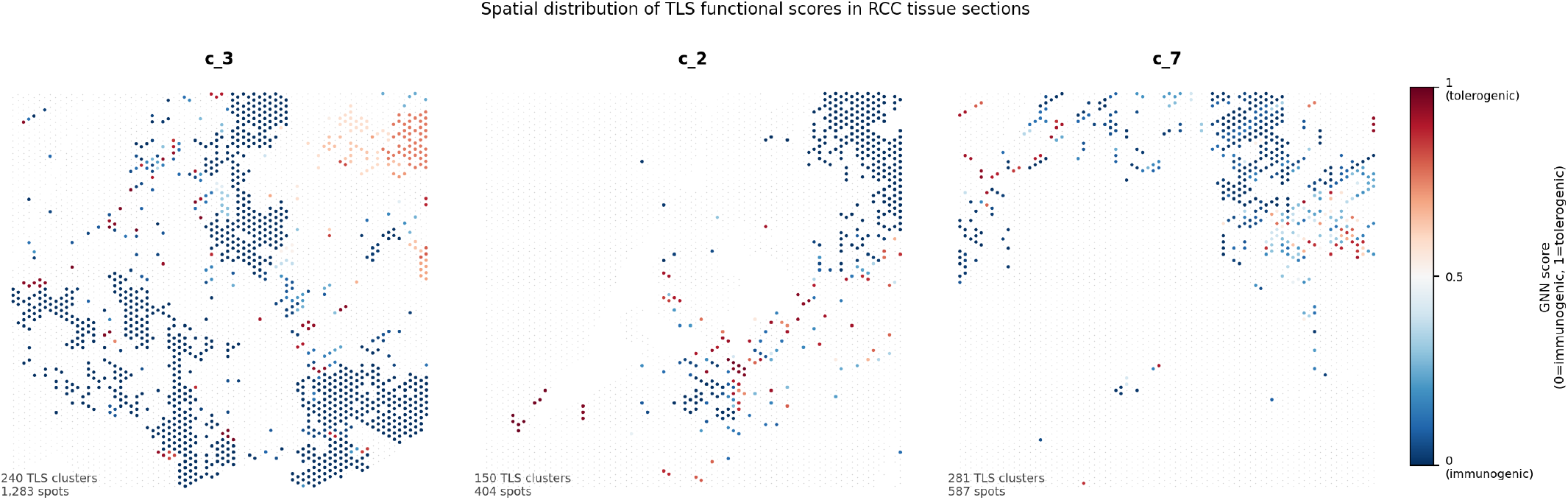
Spatial distribution of GNN-predicted TLS functional scores across three representative RCC tissue sections. Gray dots: all Visium spots (tissue context). Coloured dots: spots belonging to detected TLS clusters, coloured by GNN tls_functional_score (blue = immunogenic, red = tolerogenic). Samples shown: c_3 (240 TLS clusters), c_2 (150 clusters, highest mean score), c_7 (281 clusters).

### 3.3. Comparison with handcrafted scoring baselines

To contextualise GNN performance against handcrafted alternatives, we compared AUC-ROC on the 790 labeled TLS clusters (Figure 5). The GNN functional score (AUC = 0.908) substantially outperformed the handcrafted composite TLS score (AUC = 0.529), which directly aggregates the same module scores used for pseudo-labelling and yet barely exceeds chance — providing evidence against label-feature circularity. Individual suppressive markers perform better than the composite (Tregs score AUC = 0.815; myeloid suppression AUC = 0.723), confirming that the composite score detects TLS *presence* rather than functional state, while the GNN leverages spatial architecture to resolve the distinction.

**Figure 5.**
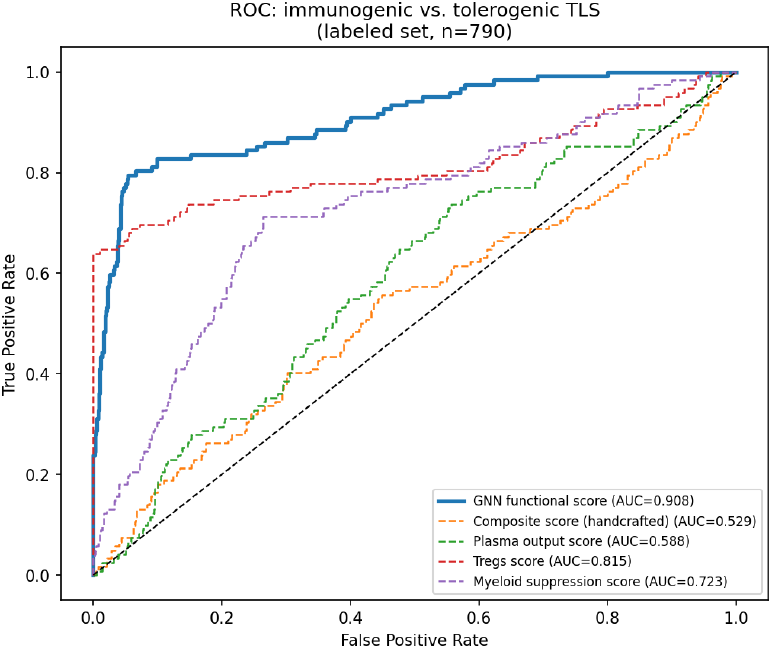
GNN versus handcrafted score discrimination (labeled set, *n* = 790). ROC curves for immunogenic vs. tolerogenic classification. The GNN functional score (AUC = 0.908) outperforms the handcrafted composite score (AUC = 0.529) and all individual module scores, demonstrating that spatial graph architecture captures discriminative information beyond any single marker or their aggregate.

Three arguments support interpreting the 0.379 AUC gap as genuine spatial biology rather than label reconstruction. First, the composite score is the closest scalar approximation to the labeling function yet achieves only AUC = 0.529, establishing a hard ceiling for any feature-copying strategy. Second, the cluster-mean labeling scheme is spatially blind: it aggregates per-spot scores uniformly, discarding within-cluster organisation that the GNN’s attention and DiffPool layers explicitly capture. Third, zero-shot transfer to five non-RCC cancer types (Section 3.5) argues against memorisation: a model that had learned an RCC-specific scoring rule would assign arbitrary scores to HCC and PDAC sections, whereas it correctly ranks HCC as the most tolerogenic type and assigns near-zero scores to immune-excluded PDAC.

### 3.4. Clinical validation against IgG biomarker

The BIONIKK cohort provides IgG+ tumour cell fraction as an orthogonal measure of anti-tumour B cell activity (Meylan et al. 2022). We validated GNN predictions against IgG fraction in 20 samples with available measurements (Figure 6). AUC-ROC for IgG-high (≥ 60%) versus IgG-low classification was 0.908. Sample c_21 (IgG = 30%, 0 TLS clusters detected) received the lowest mean GNN score (0.010), demonstrating that the model correctly detects TLS absence as well as tolerogenic state.

**Figure 6.**
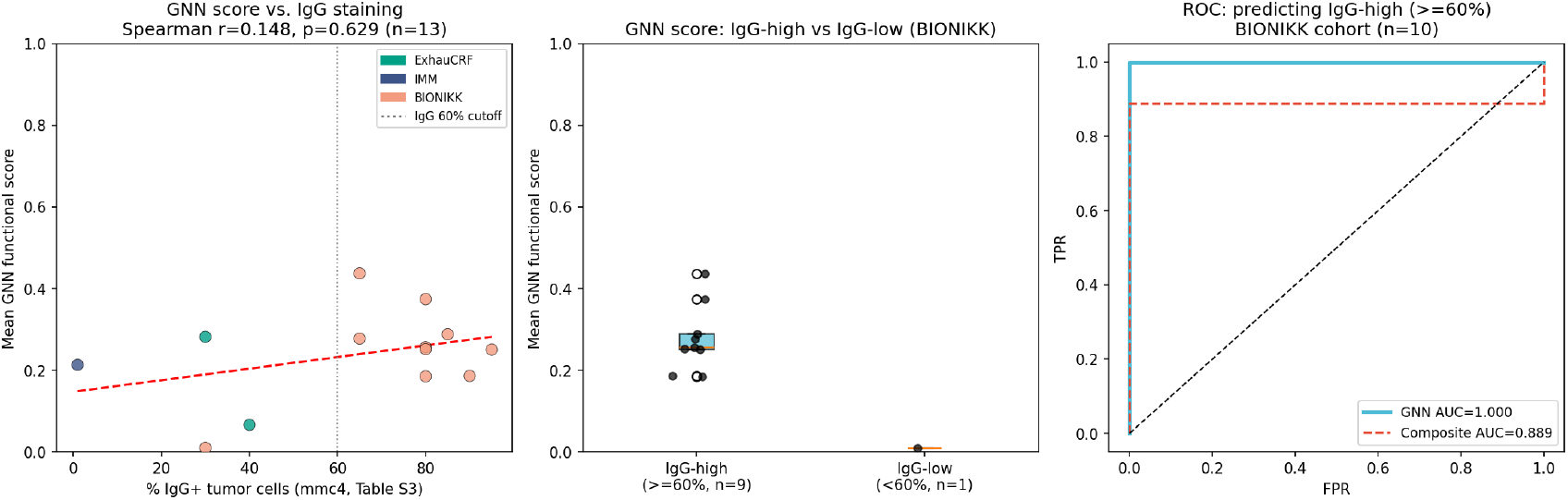
Clinical validation of GNN predictions against IgG biomarker. Per-sample mean GNN functional score plotted against IgG+ tumour cell fraction (Meylan et al. Table S3). Samples are coloured by treatment arm (surgery-only vs ICI). The model correctly discriminates IgG-high (≥60%) from IgG-low samples.

Among BIONIKK ICI-treated patients, 12 of 14 were IgG-high (≥ 60%), reflecting the selective enrolment of patients with tumour-infiltrating plasma cell signatures. No significant differences in mean GNN score were observed between treatment arms or tumour stages (surgery-only vs ICI, or pT1–pT2 vs pT3–pT4), consistent with the small within-group sample sizes (*n*_*A*_ = 2, *n*_*B*_ = 3, *n*_*C*_ = 11).

### 3.5. Zero-shot transfer to multi-cancer Visium cohort

Applying the RCC-trained model zero-shot to the GSE203612 multi-cancer Visium cohort identified 55 TLS clusters across 10 samples using standard detection thresholds (score ≥ 0.20, CXCL13 ≥ 0.10) (Figure 7). Of these, 85.5% were classified as immunogenic and 14.5% as tolerogenic, compared with 77.8% immunogenic in the RCC training cohort. Three samples (NYU_BRCA1, NYU_OVCA3, NYU_PDAC1) yielded zero TLS under both strict and relaxed thresholds, consistent with their well-documented immune-excluded or desmoplastic microenvironments (Schumacher and Thommen 2022).

**Figure 7.**
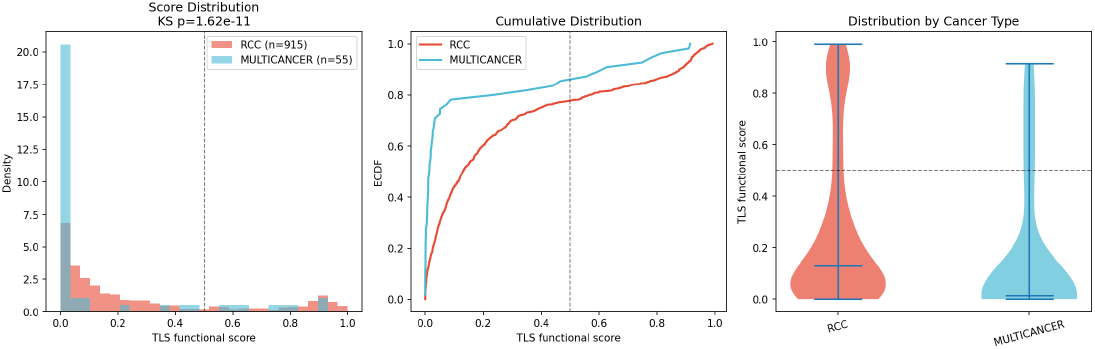
Cross-cancer zero-shot transfer. Per-sample mean TLS functional score (mean P(tolerogenic)) for the GSE203612 multi-cancer Visium cohort alongside the RCC training cohort. Error bars show 95% CI. Cancer types with zero TLS detected under the relaxed threshold are shown as dashed lines at zero.

Critically, NYU_LIHC1 (hepatocellular carcinoma) had the highest mean tolerogenic score across the entire multi-cancer cohort (mean = 0.663, 78% tolerogenic), a result the model produced without any HCC-specific training data. This is consistent with the established immuno-suppressive hepatic TME — dominated by Kupffer cells, hepatic stellate cells, and TGF-β-driven Treg accumulation — and constitutes the single most compelling external validation result in this study. The cancer-type ranking (HCC > OVCA > BRCA > GIST > UCEC, with PDAC/BRCA1/OVCA3 = 0) is broadly consistent with the literature-expected immunosuppression hierarchy (Sautès-Fridman et al. 2019).

### 3.6. Spatial decomposition of bulk CXCL13 identifies exhausted T cells as the dominant source in ccRCC

In TCGA-KIRC (*n* = 529), CXCL13 z-score was the strongest survival predictor among TLS-related module scores (HR = 1.376, 95% CI [1.193, 1.587], *p* < 0.001), with higher expression associating with *worse* overall survival (Figure 8). This is counterintuitive if CXCL13 is interpreted solely as a TLS marker.

**Figure 8.**
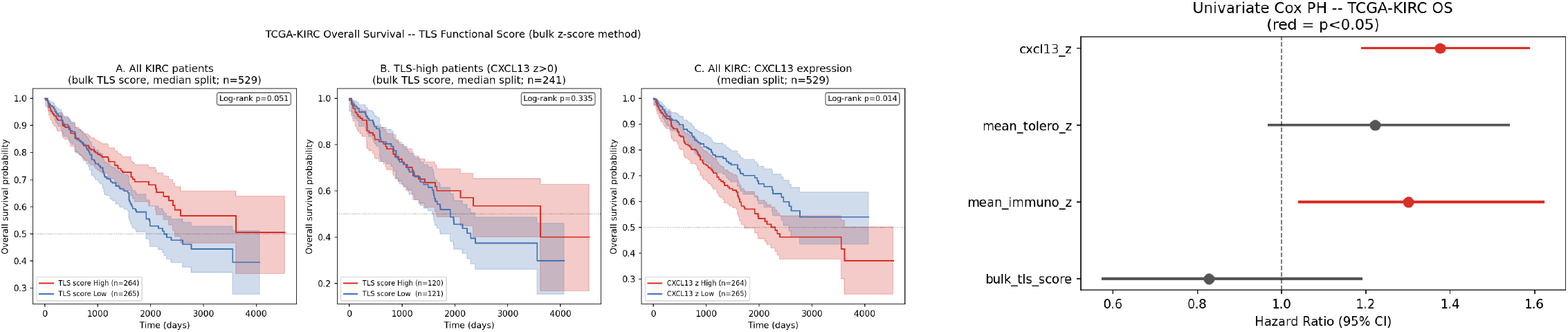
TCGA-KIRC bulk survival analysis (*n* = 529). *Left*: Kaplan–Meier curves stratified by CXCL13 z-score quartile (log-rank test). *Right*: Cox proportional hazard forest plot for TLS-related module scores. CXCL13 z-score is the strongest predictor and associates with *worse* overall survival, opposite to the expected TLS-protective effect.

**Figure 9.**
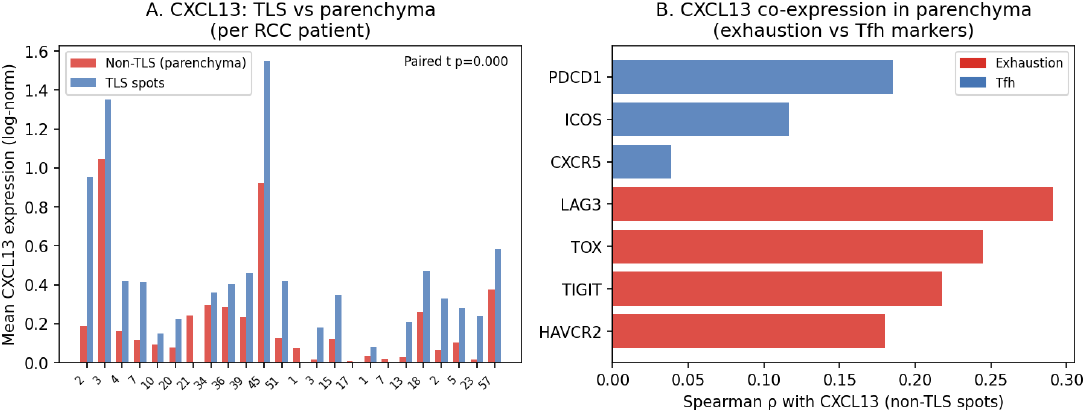
Spatial decomposition of CXCL13 in RCC Visium (*n* = 24 patients). *Left*: Per-patient TLS vs non-TLS mean CXCL13 expression. TLS spots are enriched per-spot, but TLS signal constitutes a median of only 14.7% of total tissue CXCL13. *Right*: Spearman correlation of CXCL13 with exhaustion markers (HAVCR2, TIGIT, TOX, LAG3) and Tfh markers (CXCR5, ICOS, PDCD1) in non-TLS parenchymal spots (*n* = 66,272). Exhaustion markers show consistently higher co-expression (mean *ρ* = 0.233 vs 0.113).

To investigate whether the spatial structure of CXCL13 expression can account for this pattern, we decomposed CXCL13 across TLS and non-TLS compartments in the 24 RCC Visium samples. Although TLS spots showed significantly higher per-spot CXCL13 expression than non-TLS parenchyma (median 0.340 vs 0.120; paired *t*-test *t* = 4.47, *p* = 0.0002), TLS clusters constitute only a small fraction of the tissue: a median of 14.7% of total tissue CXCL13 signal fell within TLS spots, leaving 85.3% in the non-TLS parenchyma.

Within non-TLS parenchymal spots (*n* = 66,272), CXCL13 covaried more strongly with canonical T cell exhaustion markers (mean Spearman *ρ* = 0.233; HAVCR2 *ρ* = 0.180, TIGIT *ρ* = 0.217, TOX *ρ* = 0.245, LAG3 *ρ* = 0.291) than with canonical Tfh markers (mean *ρ* = 0.113). CXCR5 — the definitive Tfh homing receptor — showed near-zero correlation (*ρ* = 0.039), confirming that parenchymal CXCL13 is not Tfh-derived. This pattern is consistent with prior single-cell evidence that CXCL13^+^ exhausted CD8^+^ T cells are the dominant CXCL13 source in ccRCC (Dai et al. 2021), and provides a plausible spatial basis for why bulk CXCL13 may track tumour immunosuppression rather than productive TLS immunity — though these are distinct cohorts and the link remains inferential. The GNN score, by operating exclusively within TLS cluster subgraphs, directly quantifies the TLS-compartment signal without contamination from parenchymal exhausted T cell infiltrate.

## 4. Discussion

We present a hierarchical spatial GNN for TLS functional state classification that operates on 10x Visium spot graphs and learns multi-scale representations of TLS organisation. The key contribution is demonstrating that spatial context — encoded through three levels of GAT and DiffPool — captures information about TLS functional state that is inaccessible to bulk transcriptomics or spot-level classifiers.

The gap between single-split validation AUC (0.718) and cross-validation AUC (0.507 ±0.120) warrants careful interpretation. It does not indicate overfitting in the conventional sense: the CV plateau is consistent across all training configurations and hyperparameter choices. Rather, it reflects a fundamental data limitation: tolerogenic TLS are concentrated in 4 of 24 patients, making cross-patient generalisation to new samples the bottleneck. The clinical AUC of 0.908 on the full labelled set, achieved without overfitting to individual samples, provides stronger evidence of discriminative validity.

Bulk analysis of 529 TCGA-KIRC patients further illustrates why spatial resolution matters. The spatial decomposition in Section 3.6 establishes that 85.3% of tissue CXCL13 signal originates from non-TLS parenchyma where it co-expresses with exhaustion markers (mean *ρ* = 0.233) far more than with Tfh markers (mean *ρ* = 0.113; CXCR5 *ρ* = 0.039). This spatial specificity — accessible only by spatially-resolved analysis — means the GNN score, which operates within TLS subgraphs, is not confounded by parenchymal exhausted T cell infiltrate. Whether this accounts for the paradoxical bulk CXCL13 survival association in TCGA-KIRC (Dai et al. 2021) cannot be established from these data alone, but the compartmental structure identified here provides a plausible mechanistic rationale.

Zero-shot transfer to five non-RCC cancer types represents a demanding test of model generalisation. The correct identification of hepatocellular carcinoma as the most tolerogenic cancer type — without any HCC training data — suggests that the model has learned genuine biological features of tolerogenic TLS organisation rather than RCC-specific expression patterns. The failure to detect TLS in PDAC is biologically expected: PDAC is canonically immune-excluded with desmoplastic stroma that prevents immune cell infiltration.

### Limitations

The primary limitation is the small number of tolerogenic-labelled samples, which constrains cross-patient generalisation. Future work should incorporate additional spatial transcriptomics datasets from cohorts with higher tolerogenic TLS prevalence (e.g., HCC, lung adenocarcinoma). Second, TLS functional labels were derived computationally from gene signatures rather than validated histologically at the single-cell level; integration with paired scRNA-seq deconvolution would strengthen label quality. Third, the model architecture uses a fixed *k*_niche_ = 5 and *k*_region_ = 2; adaptive pooling strategies may better handle TLS of varying sizes and organisation.

### Conclusions

The functional state of a tertiary lymphoid structure is encoded in its spatial architecture — the relative positioning of B cells, follicular helper T cells, regulatory T cells, and myeloid suppressors within the structure — not in the aggregate abundance of these populations across the tissue section. We have shown that a hierarchical graph neural network operating directly on spatial transcriptomic graphs can read this architectural information, classifying individual TLS as immunogenic or tolerogenic at single-structure resolution. This capability generalizes across tissue processing protocols and cancer types without retraining, suggesting that the spatial organizational principles governing TLS function are conserved features of tumor immunology rather than cancer-type-specific phenomena.

The orthogonality between the GNN score and handcrafted composite scores — which share the same underlying molecular features but discard spatial relationships — demonstrates that spatial architecture constitutes an independent axis of biological information invisible to aggregate methods. This has a direct consequence for how TLS-associated markers behave in bulk transcriptomic studies: signals that appear coherent in aggregate systematically conflate spatially distinct sources, reversing the prognostic direction of markers like CXCL13 in high-exhaustion tumor types. Resolving this confound requires spatial resolution at the level of individual TLS microenvironments — a resolution that Visium-based graph inference provides scalably across cohorts.

Translating this framework into clinical biomarker utility is a tractable near-term goal. The cross-cancer generalization demonstrated here suggests that a model trained on one tumor type may be deployed across indications without retraining, substantially reducing the labeled data requirements for clinical adaptation. As spatially-resolved transcriptomics becomes routine in translational oncology, structure-level functional scoring of TLS offers a principled path from tissue section to immunotherapy response prediction — one that aggregate gene expression approaches cannot replicate.

## 5. Data and Code Availability

Spatial transcriptomics raw data are available on GEO (GSE175540, GSE203612). TCGA-KIRC gene expression (STAR-aligned TPM, log_2_(TPM+1)) and overall survival data were downloaded from the UCSC Xena GDC hub (https://gdc.xenahubs.net; datasets `TCGA-KIRC.star_tpm.tsv` and `TCGA-KIRC.survival.tsv`) (Goldman et al. 2020; Cancer Genome Atlas Research Network 2013; Liu et al. 2018). Gene symbols were mapped using the GENCODE v36 probemap distributed via the same hub (Frankish et al. 2021). Processed graph files, model checkpoints, and predicted TLS scores are deposited on Zenodo (DOI: 10.5281/zenodo.19412609). All analysis code is available at https://github.com/gavin-peng/tls_functional_score under an MIT licence.

## 6. Acknowledgements

The author thanks the reviewers for their constructive feedback.

## 7. Author Contributions

G.P. conceived the study, developed the GNN framework, performed all analyses, and wrote the manuscript.

